# Development of a 3D *in vitro* human-sized model of cervical dysplasia to evaluate the delivery of ethyl cellulose-ethanol injection for the treatment of cervical dysplasia ablation

**DOI:** 10.1101/2024.02.20.581242

**Authors:** Ines A Cadena, Gatha Adhikari, Alyssa Almer, Molly Jenne, Ndubuisi Obasi, Nicolas F. Soria Zurita, Willie E Rochefort, Jenna L. Mueller, Kaitlin C. Fogg

## Abstract

Cervical cancer, the second leading cause of cancer-related death for women worldwide, remains a preventable yet persistent disease that disproportionately affects women in low and middle-income countries (LMICs). While existing therapies for treating cervical dysplasia are effective, they are often inaccessible in LMICs. Ethanol ablation is an alternative low-cost, accessible therapy that we previously enhanced into an ethyl cellulose (EC)-ethanol gel formulation to improve efficacy. When seeking to evaluate EC-ethanol for cervical dysplasia, we found a paucity of relevant animal models. Thus, in this study, we developed a 3D *in vitro* model of cervical dysplasia featuring a central lesion of cervical cancer cells surrounded by fibroblasts and keratinocytes to enable the evaluation of EC-ethanol and other novel therapeutics. Our GelMA-based 3D model successfully captured the architectural complexity of cervical dysplasia, showcasing cell response and high viability. The GelMA hydrogel formulation (8.7% w/v) exhibited viscoelastic properties akin to human cervical tissue. Using micro-CT imaging, we assessed EC-ethanol injection deposition in the hydrogel, revealing retention of virtually the entire injected volume near the injection site. Finally, we evaluated the EC-ethanol injection’s efficacy in eliminating cervical cancer cells. The EC-ethanol injection led to a significant decrease in cancer cell viability while preserving healthy cells in the 3D *in vitro* model. Taken together, our *in vitro* model mirrored the architecture of cervical dysplasia and demonstrated the potential of EC-ethanol for localized treatment of cervical dysplasia.

## 1 Introduction

Cervical cancer is a preventable disease, yet it remains the second leading cause of cancer-related death for women worldwide.^(1)^ Mortality is projected to rise to 400,000 annual deaths by 2030, and 90% of those deaths will be in low and middle-income countries (LMICs) if no additional action is taken.^(1)^ Cervical cancer is highly preventable through vaccination against the human papillomavirus (HPV)^(2)^ as well as screening, diagnosis, and treatment of cervical dysplasia before it progresses to cervical cancer.^(3)^ While considerable efforts have been made to increase access to vaccination, screening, and diagnosis^(4–9)^, up to 80% of women who have been diagnosed with cervical dysplasia in LMICs never receive treatment, largely due to treatment being inaccessible at the point of care.^(10)^ Thus, there is a need to develop novel therapies for cervical dysplasia that are accessible in LMICs.

In light of the above, ethanol ablation is a form of clinical tissue ablation well-suited for LMICs.^(11)^ Ethanol ablation entails directly injecting ethanol into the region of interest to induce necrosis through protein denaturation and cytoplasmic dehydration.^(12)^ It is a standard treatment option for small, unresectable liver tumors and has low complication rates.^(13)^ However, its direct injection into tissue leads to fluid leakage outside of the lesion, which decreases efficacy and can lead to side effects.^(14)^ To address this limitation, we develop a new formulation that includes a low-cost, biocompatible polymer called ethyl cellulose (EC) that forms a gel when injected into tissue. Previous studies have shown promising results in the use of EC-ethanol ablation to safely and effectively treat tumors in a variety of preclinical models, including a chemically induced hamster cheek pouch model of squamous cell carcinoma^(15)^ and subcutaneous flank tumor induced by injecting a murine breast cancer cell line.^(16)^ While we are most interested in applications in the cervix, these other models were used as no xenograft models of cervical dysplasia currently exist.^(17)^ Most recently, we scaled up to swine cervices, which are comparable in size to human cervices^.(18)^ However, currently no swine model of cervical dysplasia exists—thus, our previous study was limited to normal anatomy. Taken together, none of these widely used animal models recapitulate the microenvironment of cervical dysplasia or cervical cancer, limiting our ability to investigate how to tune EC-ethanol ablation such that it kills precancerous and cancerous cells and leaves the surrounding healthy cells alone.

Three-dimensional (3D) *in vitro* models offer potential avenues for investigating the therapeutic application of EC-ethanol ablation in the context of cervical dysplasia. However, current models, including a 3D printed model^(17)^ and an organotypic raft culture^(19)^, lack suitability for testing EC-ethanol injection due to their inability to capture cervical dysplasia’s histology and size. In a prior study, we successfully demonstrated that using gelatin methacrylate (GelMA) as the main component in a 3D *in vitro* model allowed phenotypic cell responses that captured the histology of the tumor microenvironment.^(20)^ Notably, GelMA at 8.7% w/v was observed to support cervical cancer progression.^(20)^ Despite its success in modeling cervical cancer, our 3D model reflected characteristics of metastatic cancer and is therefore unsuitable for studying cervical dysplasia. Additionally, the model’s construction in a 96-well plate restricted its size, rendering it ill-suited for testing treatments such as the EC-ethanol injection. Nevertheless, considering GelMA’s mechanical properties^(21)^, it remains a promising scaffold for developing a 3D *in vitro* model specifically tailored for cervical dysplasia with some tuning for the test of therapies involving EC-ethanol injection.

The overall goal of the present study was to develop a 3D *in vitro* model of cervical dysplasia for the application of EC-ethanol injection to cervical cancer cells in the 3D model. We developed a 3D triculture model of cervical dysplasia that is the approximate size of a human cervix, captured cell responses, and resembled the mechanical properties of human cervical tissue. We then administered EC-ethanol to the 3D model and demonstrated that we could cause necrosis of the cancer cells in the center of the model while leaving the noncancerous cells (fibroblasts and keratinocytes) on the outside of the model alive. Overall, this study demonstrated that our 3D model of cervical dysplasia captured the architecture and geometry of cervical dysplasia and could be used to evaluate the potential of EC-ethanol (and in the future other novel therapies) for localized treatment of cervical dysplasia.

## 2 Materials and Methods

### 2.1 Cell lines and reagents

Normal adult human dermal fibroblasts (NHDF-Ad) and human epidermal keratinocytes-adult (NHEK-Ad) were purchased from Lonza (CC-2511 Walkerville, MD), and SiHa cells were purchased from ATCC (ATCC® HTB-35™, Manassas, VA). All cell lines were used without additional characterization. NHDF cells were cultured in FBM fibroblast basal media (supplemented with FGM-2 SingleQuot supplements: insulin, hEGF-B, GA-1000, and FBS) (Lonza). NHEK cells were expanded in KBM gold basal medium (KGM gold supplemented with Lonza’s SingleQuot supplements: hydrocortisone, transferrin, epinephrine, GA-1000, BPE low protein-nonspecific, hEGF, and recombinant human insulin 0.5%) (Lonza). Human cervical cancer cell line SiHa from ATCC (ATCC® HTB-35™, Manassas, VA) was used without additional characterization. SiHa cells were expanded in Eagle’s Minimum Essential Medium (EMEM, ATCC), supplemented with 10% FBS (Sigma-Aldrich, St. Louis, MO) and 1% Penicillin-Streptomycin (P/S, Sigma-Aldrich). All cell lines were expanded in standard cell culture conditions (37 °C, 21% O_2_, 5% CO_2_) until passage 5. Cells were recovered at 80% confluency.

### 2.2 Cell lines and reagents

All reagents were purchased from ThermoFisher (Waltham, MA) unless stated. The cell area was evaluated by taking images of the cells over time. NHDF cells, NHEK cells, and SiHa cells were stained with fluorescent dyes. CellTracker Green CMFDA (C7025), CellTracker Blue CMF2HC (C12881), and CellTracker Red CMTPX (C34552) were used to visualize and quantify the cell area over time. Three channels (469/525nm, 628/685 nm, and 377/447 nm excitation/emission) with 9 z-stack images were taken every 12 hours for 48 hours using a Cytation 5 cell imaging multimode reader (Agilent Technologies). The distance between the z-stack is described in each experiment. Images were processed with NIH Fiji-ImageJ, and cell coverage was measured by calculating the area within the well covered by cells and dividing it by the total well area (NIH, Bethesda, MD). Cell viability was quantified by adding Calcein AM (2M, eBioscience™ Calcein AM Viability Dye, 65-0853-39, San Diego, CA) and Propidium Iodide (5 M Propidium Iodide Ready Flow™ Reagent, R37169) to the cell cultures. Following the manufacturer’s instructions, the staining solution was added on top of the wells at either 24- or 48-hours post-cultured or post-treatment. Two-channel (469/525nm and 586/647 excitation/emission) 54.8 μm z-stack and 180 μm z-stack images in a 4×4 and 4×5 montage were taken to image the 3D *in vitro* models. Images were stitched to facilitate the image analysis and quantification of live/dead cells. The mean fluorescent intensity (MFI) of live cells (stained with calcein AM) and dead cells (stained with propidium iodide) were quantified using Fiji ImageJ, and the number of cells that were alive or dead were quantified using Gen 5 software (Agilent Technologies).

### 2.3 Design of a 3D mold

We employed SolidWorks (Dassault Systèmes SolidWorks Corporation) to create a customized 3D mold that mirrors the size and histology of cervical dysplasia, featuring cancer cells on the inside and surrounded by healthy cells on the outside (**Fig. 1A**). Using polylactic acid (PLA) material with 10% infill, the mold was 3D printed at the Valley Library at Oregon State University. Subsequently, we filled the PLA mold with SORTA-Clear 40 silicone rubber (Smooth-ON, Inc., Macungie, PA 18062). Following the manufacturer’s guidelines, the mold was then cured and sterilized (**Fig. 1B**). This was done for two sizes, one with a height of 10 mm, and inner diameter of cervical dysplasia of 2 mm, and another with a height of 15 mm and inner diameter of 3mm. Both models had an outside diameter of 14 mm (**Fig. 1S)**.

**Figure 1.**
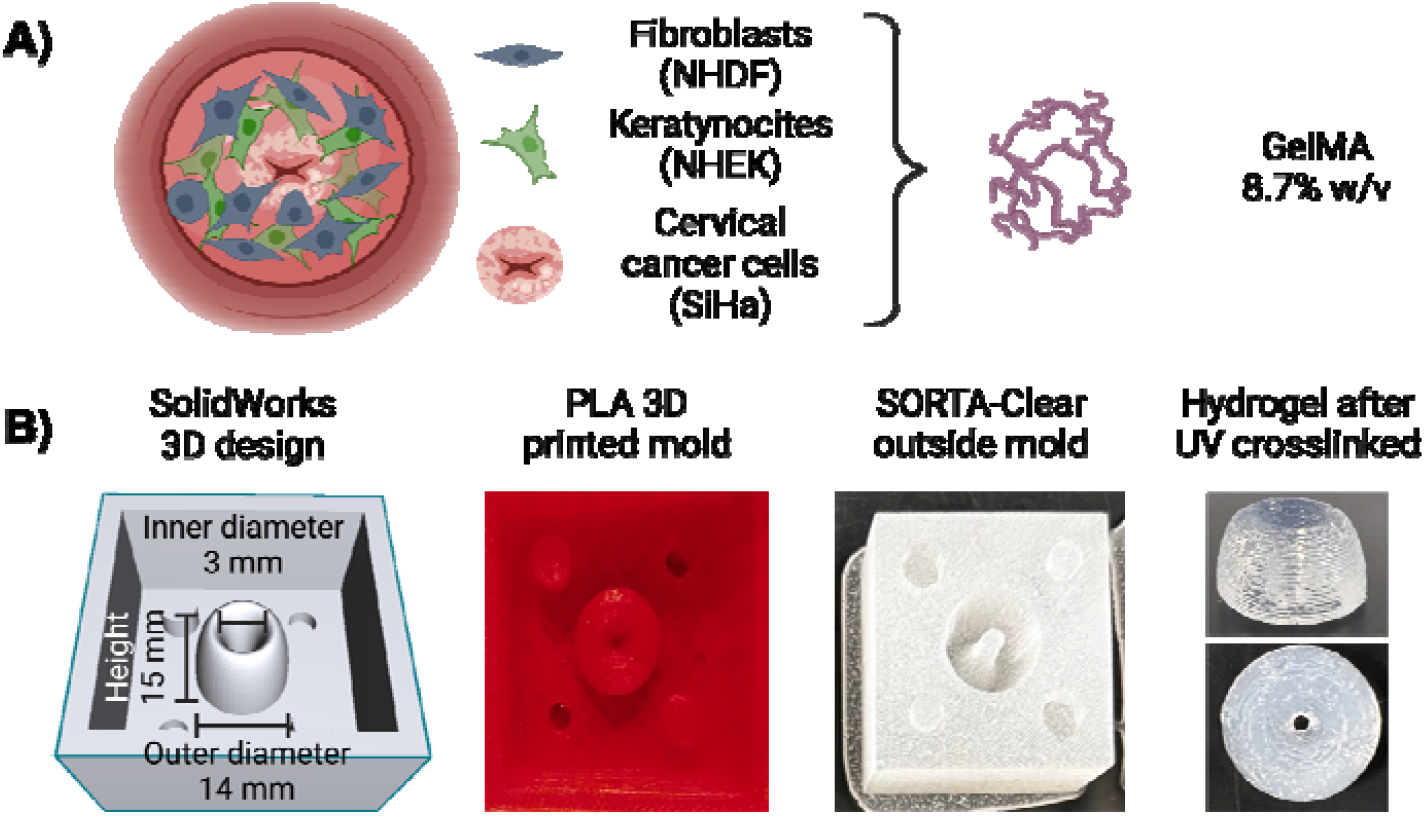
Three-dimensional (3D) mold for a tri-coculture model of cervical dysplasia. (**A**) Schematic representation of the 3D model with SiHa cells in the center encapsulated in GelMA (8.7%w/v), surrounded by NHDF cells suspended in GelMA (8.7%w/v), and covered with NHEK cells suspended in culture media. (**B**) The outside mold was created in SolidWorks, printed in PLA with a 10% infill, and built with SORTA-Clear 40 silicone rubber. Diagram created in BioRender.com

### 2.4 Assembly of 3D tri-culture model of cervical dysplasia old

We have previously reported a multilayer model of cervical cancer that used 7% w/v gelatin methacrylate (GelMA) to promote cervical cancer invasion over time.^(20)^ Following a similar protocol, we prepared the GelMA hydrogels at a final concentration of 8.7% w/v (GelMA, AdvancedBiomatrix, San Diego, CA), and we used lithium pheny-2,4,6-trimethylbenzoylphosphinate (LAP) as a photoinitiator (17 mg/mL, Advanced Biomatrix), as it has been reported to improve cell viability compared to irgacure.^(22,23)^ NHDF cells were suspended in an 8.7% w/v GelMA with LAP solution to the desired cell density for each model (**Table 1**). The final solution was pipetted into the silicone mold and UV-cured using a 405 nm light for 1 min, following the protocol specified by the manufacturer. The hydrogel was then removed from the mold and added to a black-walled 24-μ plate (IBIDI, Fitchburg, WI). Following a similar procedure, SiHa cells were suspended in a GelMA hydrogel (8.7% w/v) at the desired cell density (**Table 1**). The mixture was then pipetted in the center of the well, and UV crosslinked for 30 seconds using the same light source as before. Lastly, NHEK cells suspended in KBM-KGM culture media were added on top to simulate normal epithelium. Constructs were covered in culture media comprised of a 1:1:1 ratio of FBM-FGM-2, KBM-KGM, and EMEM. The media was changed every 24 hours, and the plate was incubated at 37 °C and 5% CO_2_ in a BioSpa live cell analysis system (Agilent Technologies, Santa Clara, CA) for 48 hours (**Fig. 2**).

**Table 1.** Cell density in the 10 mm and 15 mm constructs. The cells’ pellet was dissolved in culture media, and then cells were suspended in GelMA (8.7%w/v) hydrogels.

| Model thickness<br>(mm) | NHDF<br>(cells/well) | NHEK<br>(cells/well) | SiHa<br>(cells/well) |
| --- | --- | --- | --- |
| 10 | 53,333 | 83,160 | 26,667 |
| 15 | 133,333 | 277,200 | 106,667 |

**Figure 2.**
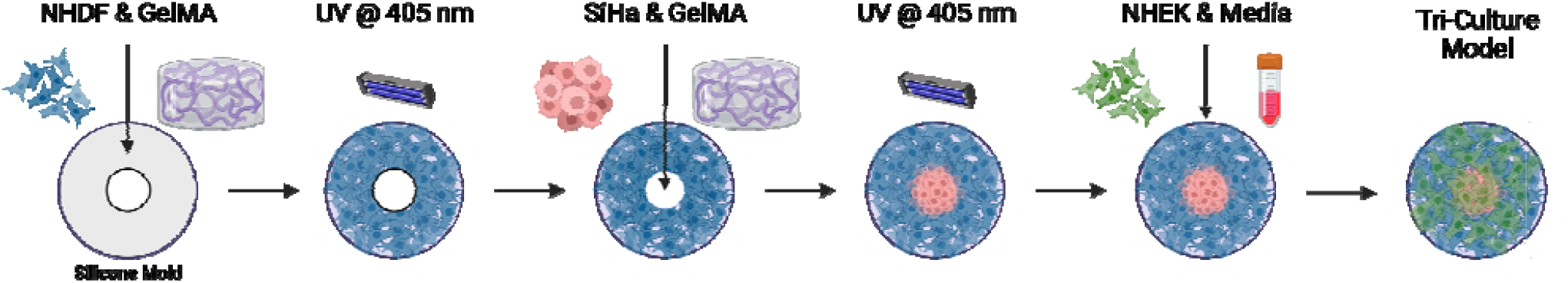
Fabrication of a 3D triculture hydrogel model for cervical dysplasia. Sche illustrating fabrication process of the 3D tri-culture model. Diagram created in BioRender.com

### 2.5 Tissue preparation and comparison of material properties between human cervix samples and GelMA hydrogel

Supplied by the University of Maryland School of Medicine (UMSOM) post-autopsy, human cervix samples were dissected to remove excess fat and residual tissue (**Fig. 3A**). For rheometric testing, the cervix was sectioned laterally to obtain samples that fit the 25mm circular plate of AR-G2 rheometer (TA Instruments, New Castle, DE). They were stored in 10% PBS and incubated at 37°C for 1 hour before experiments. We characterized the mechanical properties of our GelMA hydrogels relative to human cervical tissue using a previously described method.^(20)^ Briefly, a dynamic oscillatory shear measurement was used to evaluate the rheological properties of GelMA at 8.7% w/v (**Fig. 3B**). GelMA was kept in the incubator at 37°C before reading. Rheological characterization was evaluated using an AR-G2 rheometer (TA Instruments, New Castle, DE) equipped with a 20 mm standard steel parallel top plate with a bottom standard Peltier plate. 150 grit (120 μm particle size) sandpaper was attached to the top and bottom to prevent hydrogel from slipping. Frequency sweeps were set from 0.01 to 10 rad/s within the linear viscoelastic (LVE) region. The temperature of 37°C and a strain amplitude of 5% was kept constant during analysis. The hydrogels were allowed to equilibrate to the plate temperature for 15 min before frequency sweeps. The rheological parameters (storage moduli (G’), loss moduli (G”) and their ratio (G’/G”) were determined from these experiments.

**Figure 3.**
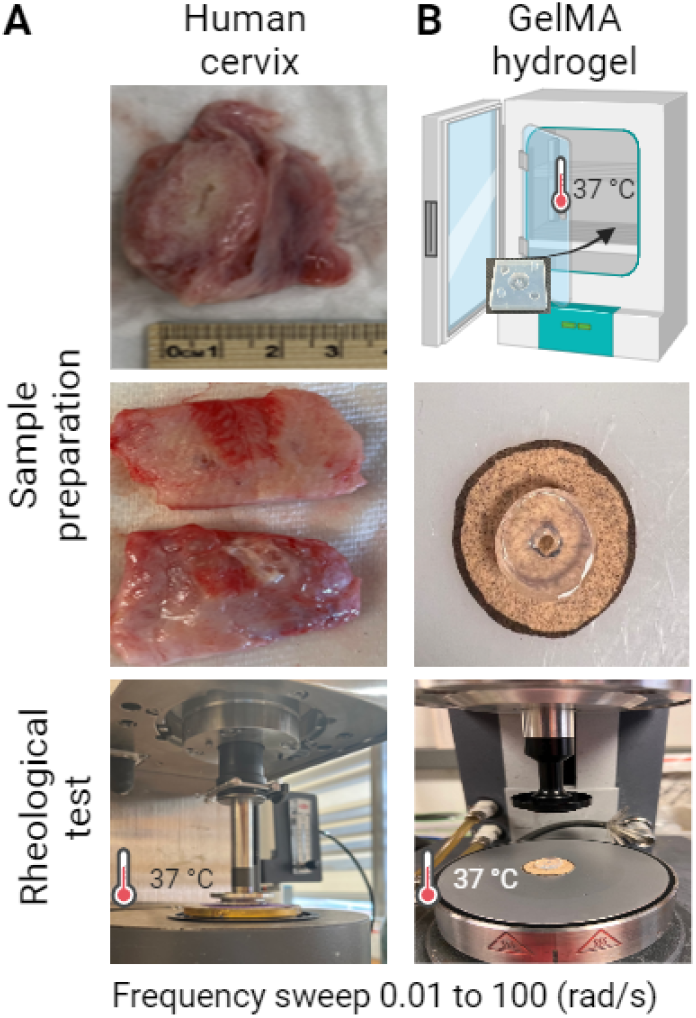
Rheological comparison between the human cervix and GelMa hydrogel. (**A**) Representation of human sample processing for rheometric test. Human samples were dissected to remove excess fat and residual tissue and cut so that they fit the 25mm circular plate (**B**). Schematic representation of sample preparation and images of GelMA (8.7%w/v) before the rheometric test. Diagram created in BioRender.com

### 2.6 EC-Ethanol formulation and contrast agent for visualization

Mixtures of ethyl cellulose (Sigma Aldrich, St. Louis, MO) and ethanol (200 proof, Koptec, King of Prussia, PA) were prepared at room temperature using a hotplate stirrer (VWR Hotplate/Stirrer). The ethyl cellulose and ethanol mixture (EC-ethanol) were continuously stirred until the ethyl cellulose was completely dissolved. In all experiments, EC concentrations of 12% (EC to ethanol, % weight: volume) were used, as previous studies indicated significant increases in distribution volume for ≥6% EC(24). To contrast the CT image between the injectate and surrounding cell construct, iohexol (Omnipaque, GE Healthcare, Chicago IL) was added to EC-ethanol. Iohexol was chosen for its high solubility in ethanol. The EC-ethanol-iohexol solutions were injected into the cell constructs and imaged using micro-CT (Bruker SkyScan 1276, Billerica, MA) as previously described.^(24)^ A 40 mg/mL concentration of iohexol was used for all experiments based on previous study results that maximized the signal-to-background ratio.

### 2.7 Micro-CT Imaging

The Bruker SkyScan 1276 micro-CT was employed for imaging to visualize the EC-ethanol distribution. Samples were positioned upright during imaging, with the needle insertion site oriented at the 12 o’clock position. The images had a pixel size of 81.072 μm, a field of view of 9.31 × 158.75 cm, and were acquired in full rotation (360□) in a 504 × 336 matrix. After image acquisition and reconstruction, post-alignment correction and appropriate ring correction were applied. EC-ethanol distribution volume was then quantified within the hydrogel by loading the images onto 3Dslicer software (Kitware, Clifton Park, NY) and maximum entropy auto-thresholding was applied to the data to segment EC-ethanol-iohexol.^(25)^

### 2.8 EC-ethanol injection in 3D triculture hydrogel model

We evaluated the effect of an EC-ethanol injection (EC to ethanol, weight to weight, i.e., 12% EC to 88% pure ethanol) on cell viability and cancer cell area in the 3D *in vitro* model. Using the 15 mm thickness construct, we co-cultured the three cell lines for 24 hours at 37 °C and 5% CO_2_ in the BioSpa. Then, 50 μL of 12% EC-ethanol was injected into the 3D *in vitro model* at a needle depth of 7 mm using a 22 G syringe. Culture media was changed every 24 hours, and images were taken using the same procedure described above. Two-channel and three-channel 180 μm z-stack images were taken every 12 hours for 48 hours post-injection to evaluate changes in cell viability and cancer cell area over time. Finally, to examine the accurate placement of the ethyl-cellulose drug within the 3D *in vitro* model, we employed a bright-field imaging using a Cytation 5 cell imaging multimode reader (Agilent Technologies). Images were stitched and z-projected prior to analysis of MFI and cancer cell area. As a control for cell viability, the 3D models were submerged in ethanol (99% v/v) for 48 hours.

### 2.9 Statistical analysis

quartile range (if an approximate normal distribution was not achieved after an appropriate transformation). Statistical analysis was performed using a two-way analysis of variance (ANOVA) with Tukey correction for multiple comparisons or one-way ANOVA when appropriate. The same analysis was used to compare human cervices samples to the GelMA hydrogels. All statistical analyses were performed using Prism 8.2.1 software (GraphPad, San Diego, CA); *p* values less than 0.05 were considered statistically significant. Replicates are specified in each experiment, and asterisks denote significance.

## 3 Results

### 3.1 Cell viability and cell coverage in 3D in vitro models of cervical dysplasia

We quantified the number of live versus dead cells in the 3D model during 48 hours of culture and saw that, over time, cell viability was maintained throughout the 3D model (**Fig. 4**). There were no significant differences in cell viability as a function of z-position. However, there was a slight decrease in the number of viable cells in the bottom layer at 24 hours (**Fig. 4C**). The 15 mm construct followed similar trends (**Fig.2S**).

**Figure 4.**
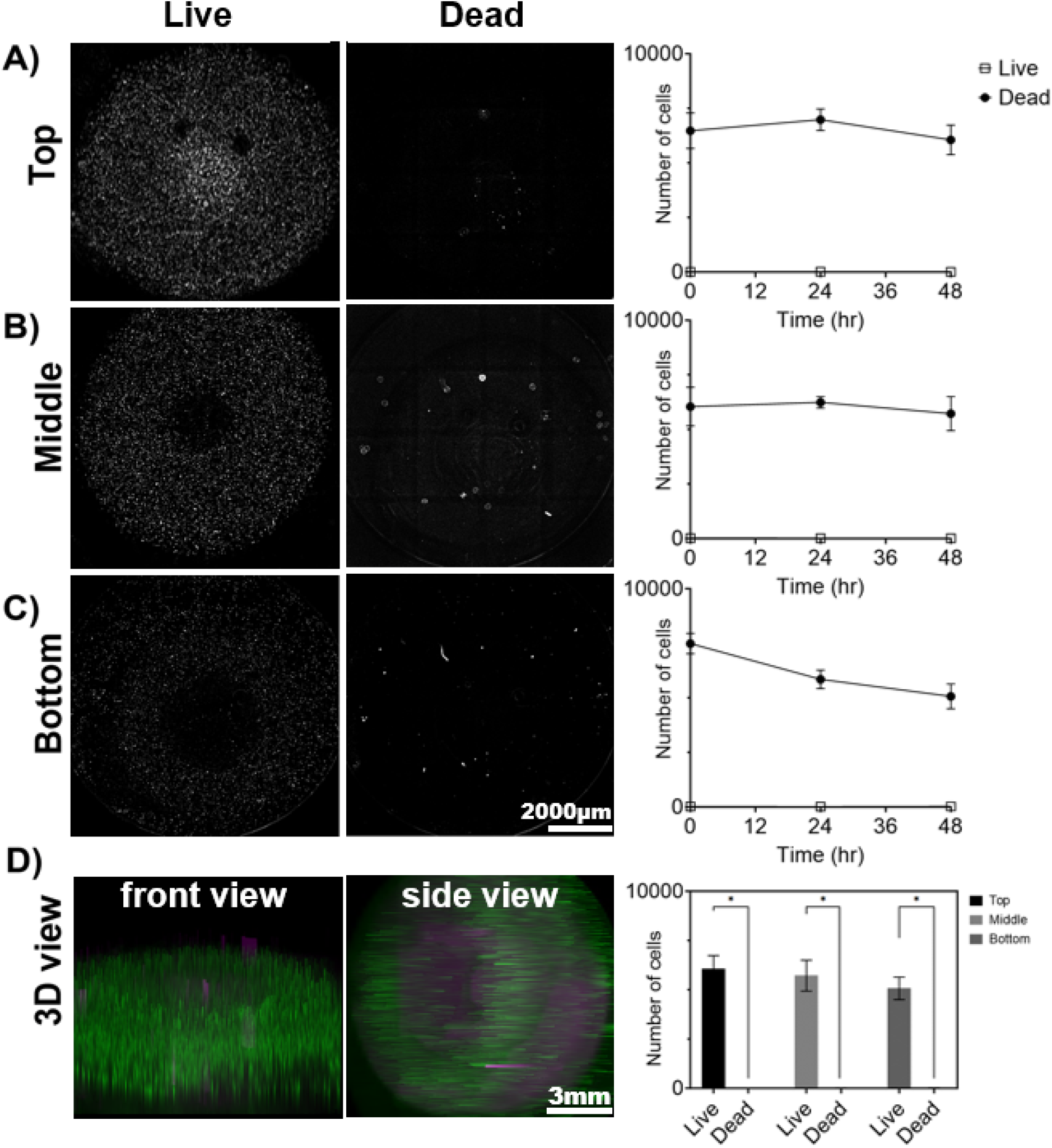
Cell viability in the 3D model for cervical dysplasia. Cell viability was measured by quantifying the number of live and dead cells in the (**A**) top, (**B**) middle, and (**C**) bottom layers of the construct at 0, 24, and 48 hours of culturing. Cells were stained with a Calcein and Propidium Iodide solution at every time point. (**D**) A 3D view of the live cells (green) and dead cells (magenta) was created using Fiji ImageJ. Images represent 48 hours after culturing. A montage of 4×4 (rows x columns) with 9 z-stacks was taken to capture the 3D model. The sample thickness was 493.2 μm, and the scale bar was set at 2000 μm. Images were created using Gen5 V3.14 software. The data represent the mean ± SEM (n = 4). \**p* < 0.05, analyzed using 2-Way ANOVA with Tukey post-test.

Our 3D model captured the architecture and geometry of cervical dysplasia, with cancer cells in the center and healthy fibroblasts and keratinocytes surrounding them. We evaluated changes in cell area over time (**Fig. 5, Fig. 2S**). At time 0 hours, cell coverage was highest for SiHa cells on the inside, and the outside had higher coverage of NHDF cells and NHEK cells. After 48 hours of culturing, there was an increase in NHEK coverage on the inside, and NHDF cells moved slightly to the inside. However, at 48 hours SiHa cells remained the predominant cell type in the inside region, and NHDF cells were predominant in the outside region (**Fig. 5B**). NHEK and NHDF cells increased their cell coverage over time, while SiHa cells maintained constant cell coverage throughout the construct after 48 hours (**Fig. 5C**). The 15 mm model was demonstrated the same trends (**Fig. 2S**). Overall, these results showed that after 48 hours of culturing, the cells were viable throughout the model and mimicked the size and architecture of cervical dysplasia.

**Figure 5.**
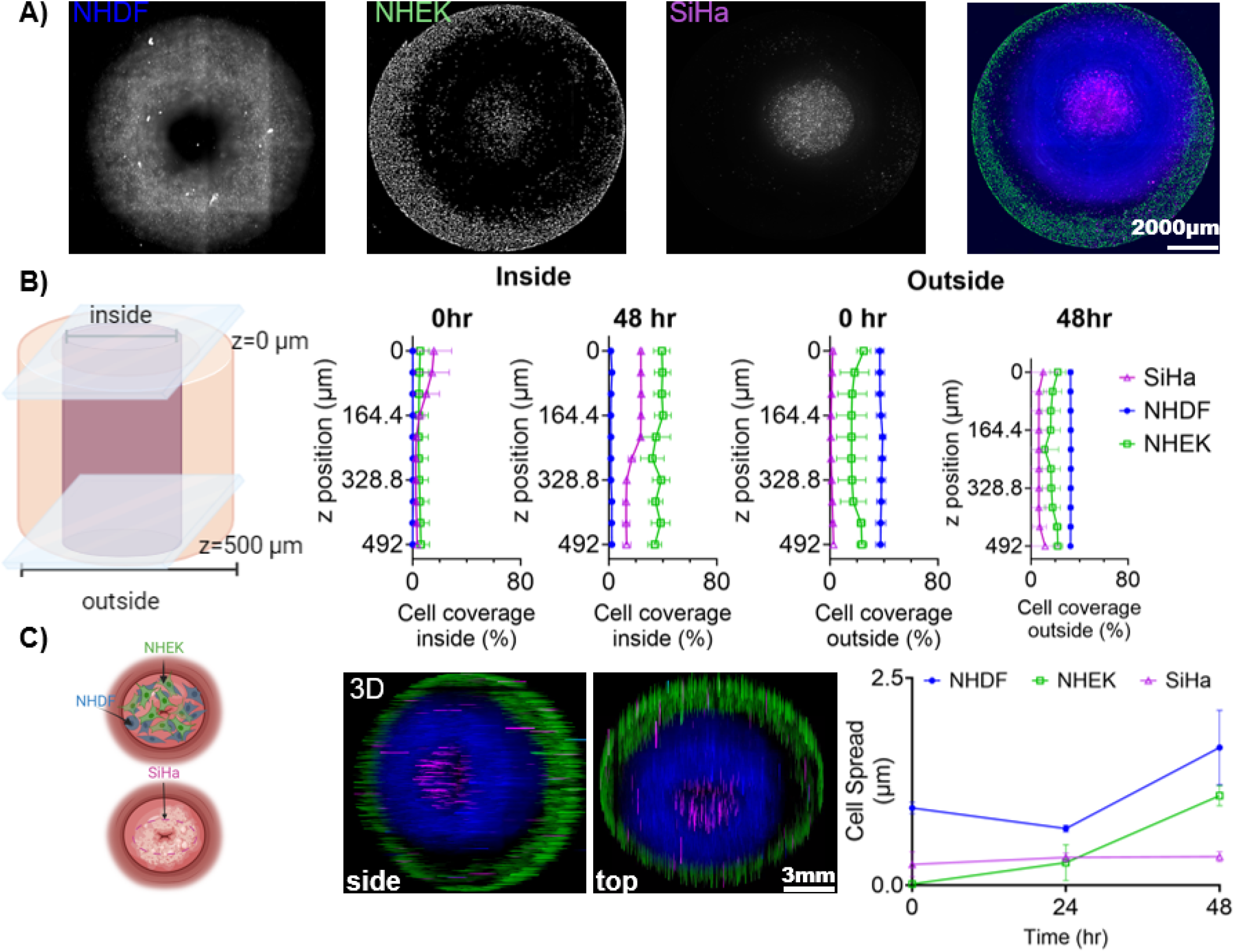
Phenotypic Cell Response in Cervical Dysplasia 3D in vitro Model. (**A**) Representative z-projection of the images of the three cell lines (NHDF, NHEK, and SiHa) after 48 hours of culturing in the 3D model. (**B**) Representation of the outer and inner regions of the 3D model, with cell coverage quantification along the z-axis. (**C**) Schematic representation of the 3D model with a view of the cells after 48 hours of co-culture. Cell spread was measured as the area covered by each cell line. Data was normalized to the time 0 and showed as a fraction. The 3D hydrogel model was cultured in a 24-well ibidi plate for 48 hours, with media changed every 24 hours. A montage of 4×4 with 54.8 z-stacks was taken to capture the 3D model. Images were created and processed with Gen5 V3.14 software. 3D z-projections were generated using Fiji ImageJ (showing the side and bottom view). Cell area was measured using Fiji ImageJ. Data represent the mean ± SD (n = 3). Schemes created in BioRender.com

### 3.2 Comparison of material properties between the human cervix and GelMA hydrogels

To ensure that the biophysical properties of our 3D model matched the tissue we were trying to recapitulate, we characterized the mechanical properties of our GelMA hydrogels relative to human cervical tissue (**Fig. 6**). Our 8.7% w/v GelMA had very similar storage moduli compared to human cervices, indicating that it captured the elastic contribution of cervical tissue’s resistance to deformation (**Fig. 6C**). The loss moduli of the GelMA were lower, suggesting that we may need to add in additional ECM proteins or explore a wider range of GelMA hydrogel formulations to capture elastic and viscous contributions (**Fig. 6D**). Our results had a higher G’/G” value for the GelMa hydrogels, suggesting its greater elasticity than the human cervical tissue (**Fig. 6E**).

**Figure 6.**
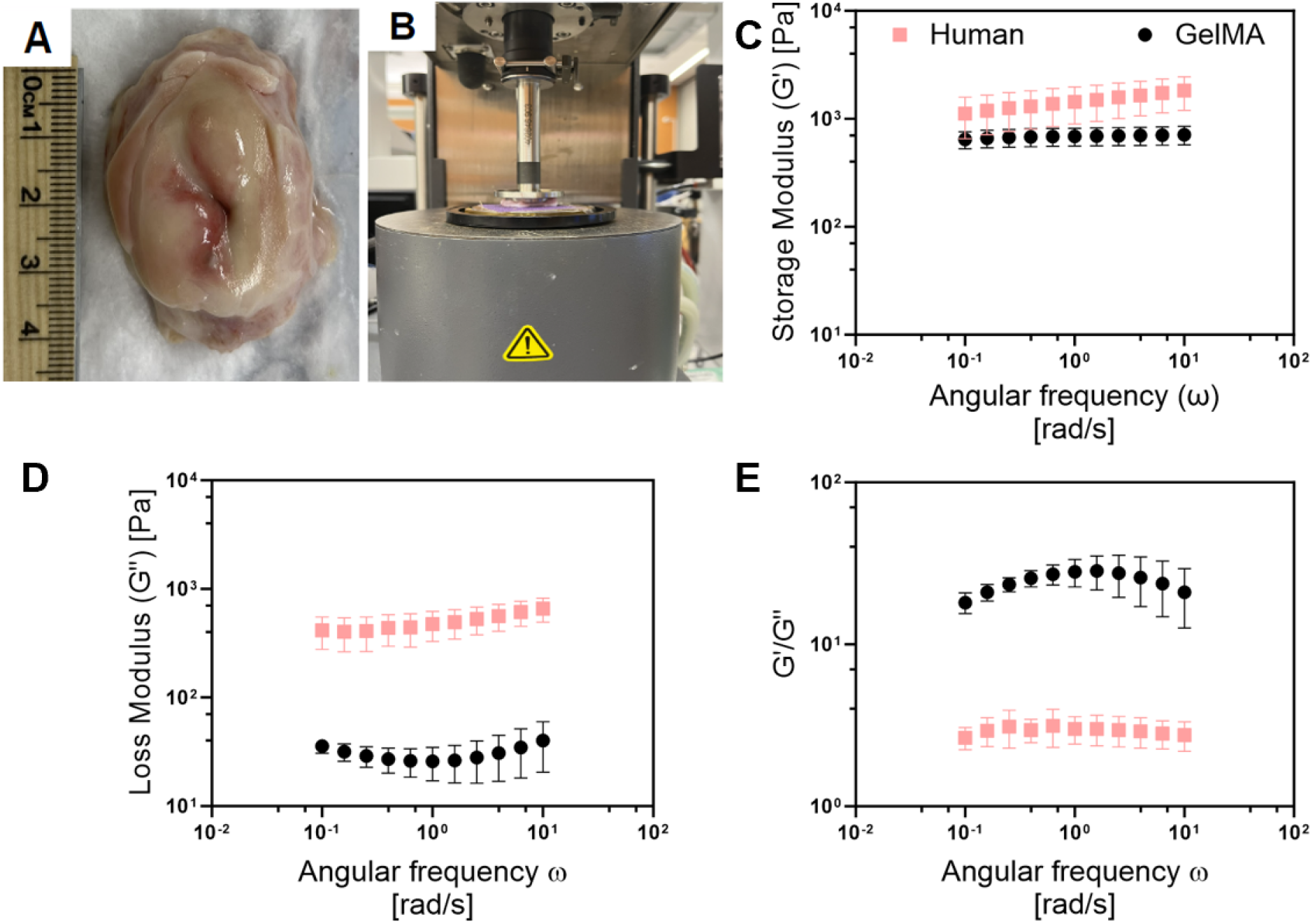
Comparison of the viscoelastic properties between the human cervix and GelMA. **(A)** Picture of human cervix obtained after autopsy from UMSOM. **(B)** Set up for rheometric testing with AR-G2 rheometer. Comparison of **(C)** storage and **(D)** loss modulus, and (**E**) G’/G” ratio of human cervices and GelMA hydrogel (8.7% w/v). Frequency sweeps were conducted from 0.1 to 100 rad/sec within the linear viscoelastic (LVE) region at a constant strain amplitude of 5%, and temperature of 37°C. Data represent the mean ± SD (n = 5).

### 3.3 Micro-CT imaging to visualize distribution of the injectate

GelMA hydrogels (8.7% w/v) were injected with EC-ethanol-iohexol (using injection parameters optimized in a previous study).^(24)^ Using Micro-CT imaging, we evaluated if iohexol (**Fig. 7B**) correlated with where the EC-ethanol injectate went, which could be visualized by its white color against the translucent GelMA background (once the GelMA was sectioned in half) (**Fig. 7A**). We demonstrated that there were no statistical differences between the two visualization methods (**Fig. 7D**), which in the future will enable us to track where the gel forms in our intact 3D *in vitro* models without the need for sectioning. Moreover, our results indicated that for injected volumes of 50 uL, virtually the entire injected volume was retained near the site of injection (**Fig. 7C**). This not only facilitates comprehensive visualization but also empowers precise quantitative analysis of the distribution of EC-ethanol in our *in vitro* constructs.

**Figure 7.**
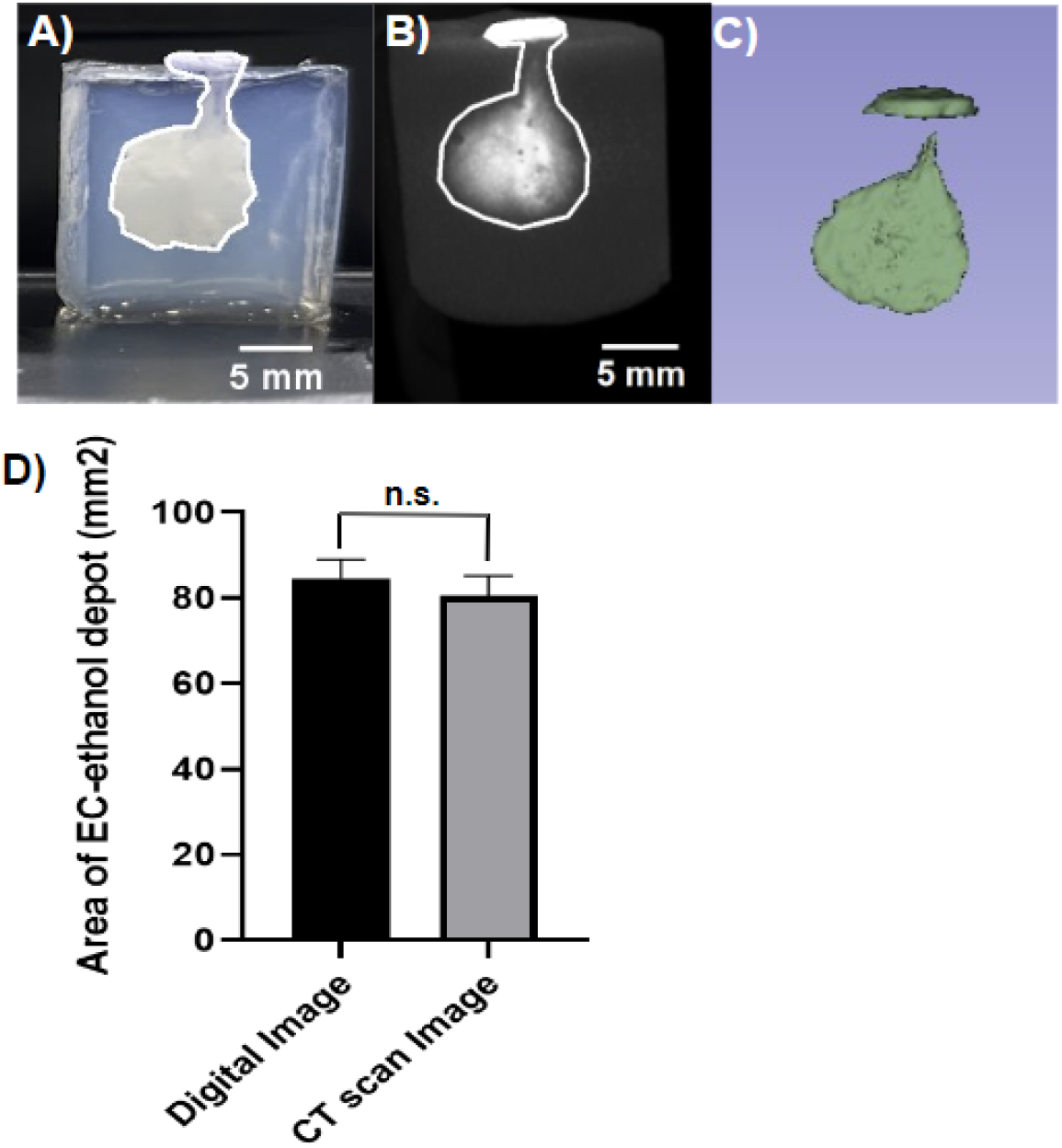
CT imaging to visualize distribution volume achieved with EC-Ethanol injection. Representative images of 12% EC-Ethanol injections on 15 mm tall 8.7 % 8.7% GelMA hydrogels with Iohexol as a contrast agent. (**A**) Digital image of the EC-Ethanol gel depot with area of 86.93 mm^2^. (**B**) Image acquired from CT imaging with area of 83.36 mm^2^. (**C**) Coin shaped 3D projection of the depot acquired using 3D slicer. The following parameters were used: 22G -25 mm-long beveled needles, 7 mm needle insertion depth, manual injection, 12% EC-ethanol, 80 mg/mL Iohexol. Images are shown as a side view. Scale bar = 5 mm. (**D**) Quantification of the area calculated from digital image vs CT scan image. Data represent the mean ±SD. No statically significant differences were observed

### 3.4 Effect of ethyl cellulose-ethanol injection on cell viability and cancer area in 3D *in vitro* model

In our 3D *in vitro* model, we evaluated the efficacy of the EC-ethanol injection in reducing the cell viability of the cancer cells and preserving the viability of the surrounding healthy cells (**Fig. 8**). We injected 50 μL of EC-ethanol and evaluated cell viability in the 3D model compared to positive and negative controls. The 3D models cultured in media had viable cells throughout the construct with very few dead cells (**Fig. 8A**). In contrast, the 3D models submerged in ethanol had only dead cells (**Fig. 8B**). Finally, the 3D models injected with the EC-ethanol had higher cell viability outside the EC-injection and higher cell death in the injection area (**Fig. 8C**).

**Figure 8.**
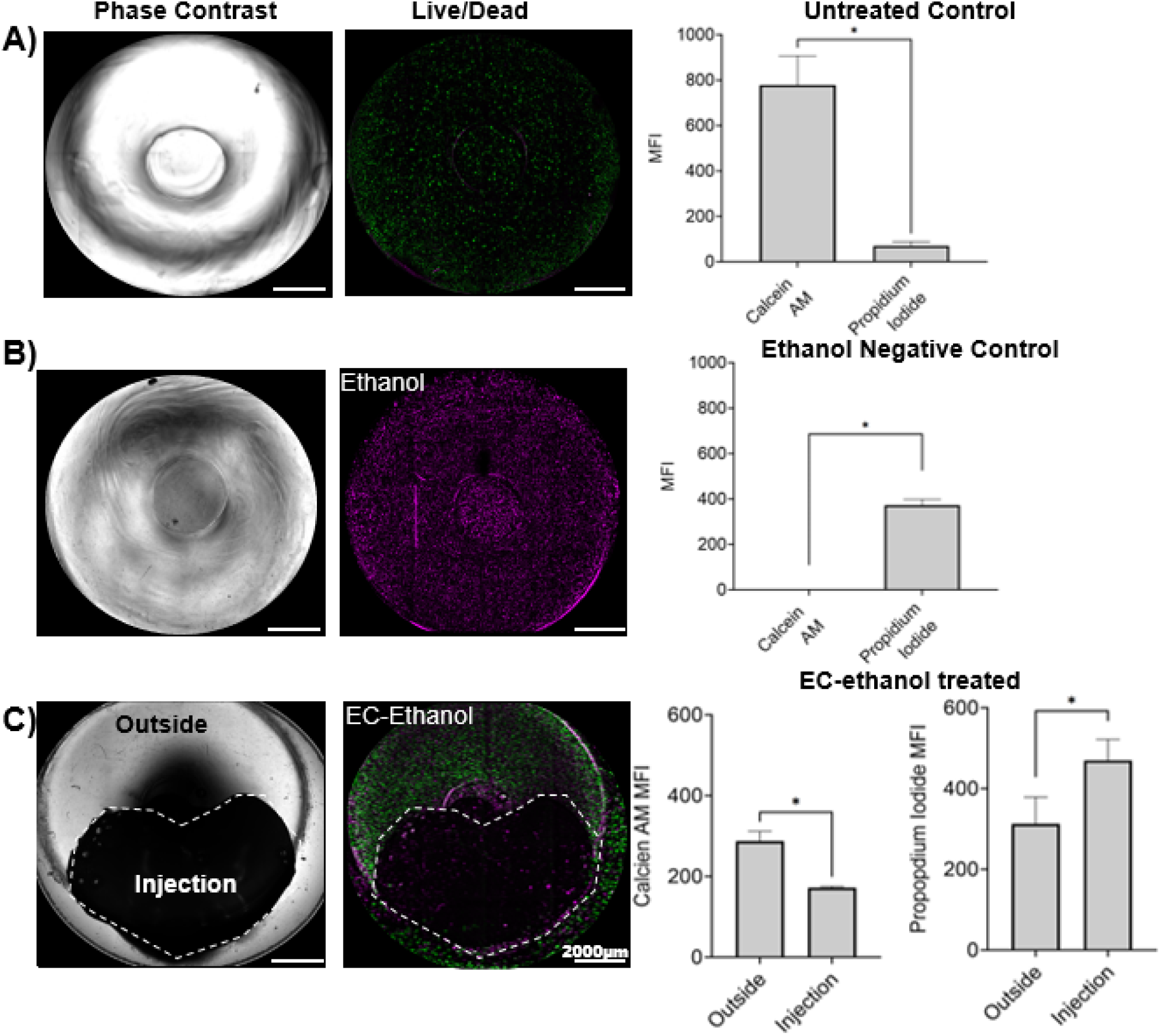
Effect of EC-ethanol on cell viability. 3D *in vitro* models **(A)** untreated and treated with (**B**) ethanol (99% v/v), and with (**C**) EC-ethanol injection. Representative images in bright field and with the fluorescent dies representing the live cells in green and the dead cells in magenta. Data represent at 48 hours post treatment. As control we culture the 3D in vitro model in (**A**) culture media, and (**B**) in ethanol for 48 hours. The mean fluorescent intensity (MFI) of live cells (stained with calcein AM) and dead cell (stained with propidium iodide) was quantified using Fiji ImageJ. The scale bar was set at 2000 μm. Images were created and processed with Gen5 V3.14 software. 3D z-projections were generated using Fiji ImageJ. * *p* <0.05, compared with the mean of each group. Paired t-test. Data represent the mean ± SD (n = 3).

We then evaluated the ability of the EC-ethanol injection to reduce the cancer cell area, as the goal in patients would be to eliminate the spread of cancer cells to the surrounding healthy tissue (**Fig. 9**). NHDF, NHEK and SiHa cells were stained with cell tracker dies and cultured in the 3D model for 24 hours prior to injection; as control, we cultured the 3D models in culture media (**Fig. 9A**). After injection there was a significant decrease in cancer cell area in the 3D models treated with the EC-ethanol injection (**Fig. 9B**), and 48 hours post-injection the cancer cells were completely eliminated. In contrast, the cancer area increased in the untreated control group.

**Figure 9.**
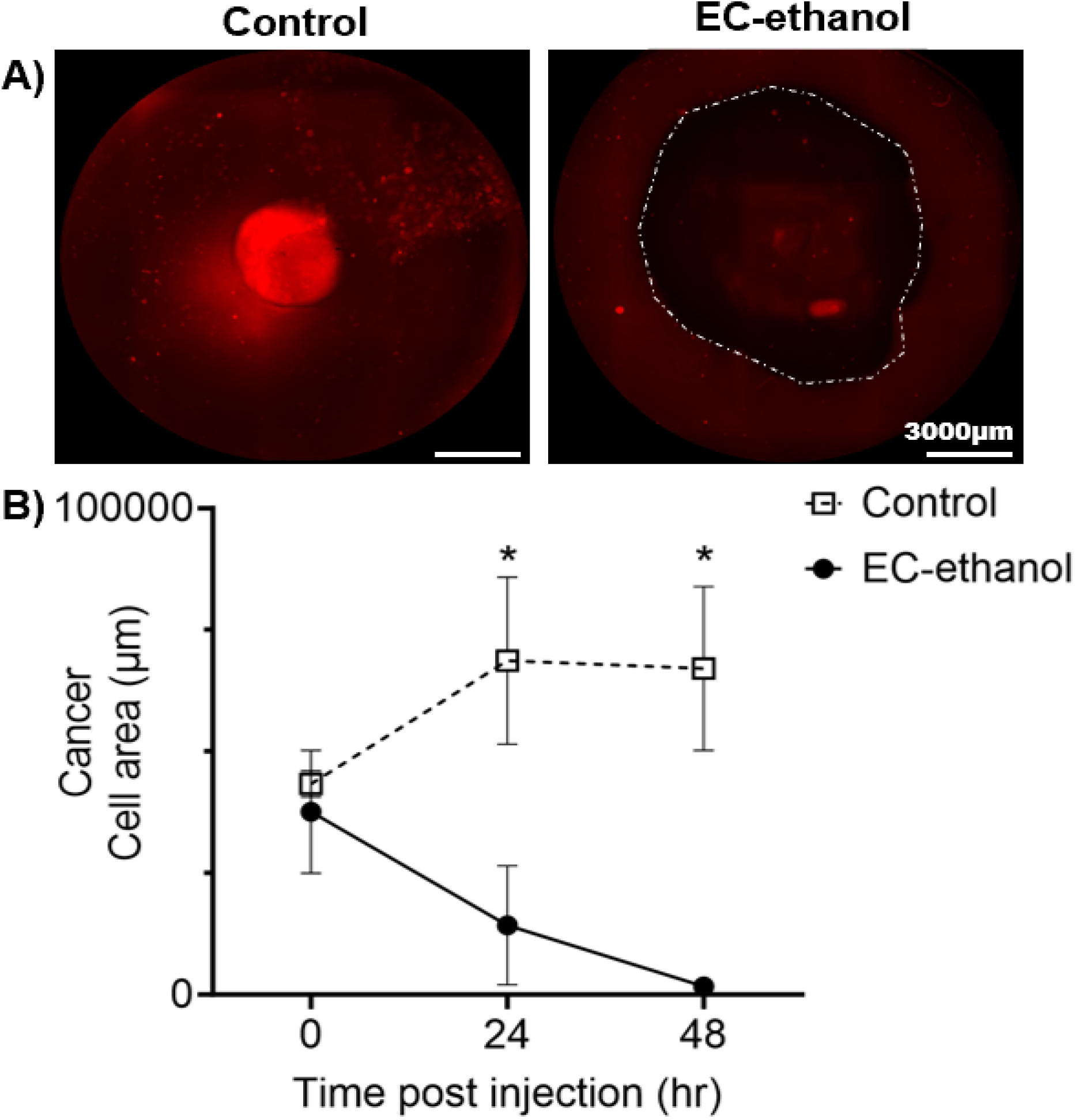
Effect of EC-ethanol on cell area. **(A)** Representative images of cancer cells cultured in 3D *in vitro* model at 48 hours with and without EC-ethanol injection. 2D z-projection full construct. **(B)** Cancer cell area at all time points pre (time 0) and post injection (24h and 48h). Control group had only culture media. Cancer cell area was quantified using Fiji ImageJ. The scale bar was set at 3000μm. Images were created and processed with Gen5 V3.14 software. 2D z-projection was created using Fiji ImageJ. * *p* <0.05, compared to the EC-ethanol group. Two-way ANOVA with Tukey post-test. Data represent the mean ± SEM (n = 4).

These findings highlight the capacity of our model to provide a more comprehensive understanding of cellular reactions to the EC-ethanol injection and its efficacy in treating disease while preserving the surrounding healthy tissue.

## 4 Discussion

Existing animal models fail to replicate the intricate microenvironment characteristic of cervical dysplasia accurately. Recognizing this significant knowledge gap, we developed a 3D *in vitro* model to mimic the architectural complexity of cervical dysplasia, featuring a central region with cervical cancer cells surrounded by healthy fibroblasts and keratinocytes. Our innovative model aimed to recapitulate the microenvironment of cervical lesions for evaluating the efficacy of ethyl cellulose (EC)-ethanol injection in mitigating cancer cell viability while preserving healthy tissue. Our 3D *in vitro* model is the first co-culture system for cervical dysplasia in the literature and is the first to use an *in vitro* model to demonstrate EC-ethanol efficacy.

We selected GelMA as the supportive matrix for human fibroblast cells and cervical cancer cells due to its biocompatibility and wide application in tissue engineering.^(26)^ We conducted a rheological characterization of the GelMA hydrogel formulation used in our model and compared its mechanical properties with human cervical tissue. We confirmed that the GelMA hydrogel-based model demonstrated compatibility with human cervical tissue in terms of mechanical properties, and their values range in similar magnitudes to those observed in the literature.^(27)^ Our results showed an average G’ of 1,453 Pa and a G” of 494 Pa in the human cervix. Meanwhile, the GelMA hydrogel had a G’ of 686 Pa and a G” of 30 Pa. Differences in the mechanical properties between our material and the human cervix are likely due to the ECM-components constituting the cervical tissue.^(28)^ However, the GelMA hydrogel has enough flexibility in its mechanical properties that allows the incorporation of more ECM components to approximate the human cervix better, a focus of future work. ^(20,29)^

We also showed that the GelMA-hydrogel model had high cell viability and promoted cell growth over 48 hours. GelMA has proven to have good transport of nutrients and oxygen that enables cell response over time.^(30,31)^ A recent model showed the encapsulation of human pancreatic cells and stromal fibroblasts in GelMA (8% w/v) with the photoinitiator LAP could mimic cell-ECM interactions and is proven to have a good diffusion of nutrients and oxygen, which allowed the cells to proliferate in the GelMA beads.^(32)^ Using the GelMA hydrogel, we were able to increase the thickness of our first model and still have a higher number of live cells compared to dead cells. Moreover, in this study, we observed that cell viability and function were maintained in both models (10 mm and 15 mm), with the presence of healthy cells mostly on the outside and cancer cells on the inside. Over time, there was a slight spread of the cancer cells to the outside and an increase in cell area for the keratinocytes that were found in all the regions at 48 hours.

To demonstrate a potential application of our 3D *in vitro* model, we evaluated the distribution area achieved with our EC-ethanol gel (through microCT imaging) and the resulting region of cell viability and cancer cell area post-injection. A critical finding of this work is that cell viability drastically decreased in the injection area, while the outer region remained unaffected, suggesting the targeted, localized impact of EC-ethanol. As previously reported, studies suggested that a concentration higher than 6% of EC-ethanol gave a localized injection with minimal leakage and highlighted the potential of EC-ethanol in precisely targeting diseased areas.^(33)^ The complement analysis on cell viability and cell area gave a better understanding of cell response to the EC-ethanol treatment in comparison with previous studies that have reported only the effect of the volume distribution of EC-ethanol.^(33,34)^ These findings highlight the nuanced and comprehensive understanding that our model offers regarding cellular reactions to EC-ethanol, emphasizing its promise as an ideal model in which to study the relationship between EC-ethanol distribution and cellular response. Further studies with the inclusion of ECM components specific for cervical dysplasia could delve into the molecular mechanisms underpinning these observed effects, further advancing our understanding of how to best deliver EC-ethanol to treat cervical dysplasia.

## 5 Conclusion

We developed a 3D *in vitro* model of cervical dysplasia that demonstrated the successful elimination of cervical dysplasia following EC-ethanol injection while leaving the surrounding healthy cells intact. These results show the promise of using our 3D *in vitro* model to study localized interventions in cervical dysplasia, and the insights gained from EC-ethanol ablation in the 3D *in vitro* model will inform its future *in vivo* applications. While our initial findings are promising for treating lesions, critical questions remain regarding optimal delivery parameters, particularly as lesions progress from low-grade to high-grade dysplasia to early invasive disease. These parameters are pivotal for the effective delivery of EC-ethanol into the cervix, ensuring comprehensive coverage of precancerous lesions and early invasive disease within deeper tissue layers. Further investigation of these delivery parameters is essential for the successful translation of EC-ethanol and similar therapeutic solutions for treating cervical dysplasia in LMICs.

## 6 Conflict of Interest

The authors declare that the research was conducted in the absence of any commercial or financial relationships that could be construed as a potential conflict of interest.

## 7 Author Contributions

I.C., G.A., K.F., and J.M. conceived the study. A.A., M.J., N.O., I.C, and G.A. performed the studies and collect data. N.S. and W.R designed the mold for the triculture model. I.C. and G.A. analysed the data. I.C., G.A., K.F., and J.M. wrote and edit the manuscript.

## 8 Data Availability Statement

All the images are available on the Fogg Lab GitHub (https://github.com/fogg-lab/Cervical-Dysplasia). The datasets generated during and analyzed during the current study are available from the corresponding author on reasonable request.

## Supplemental Figures

**Supplementary Figure 1.**
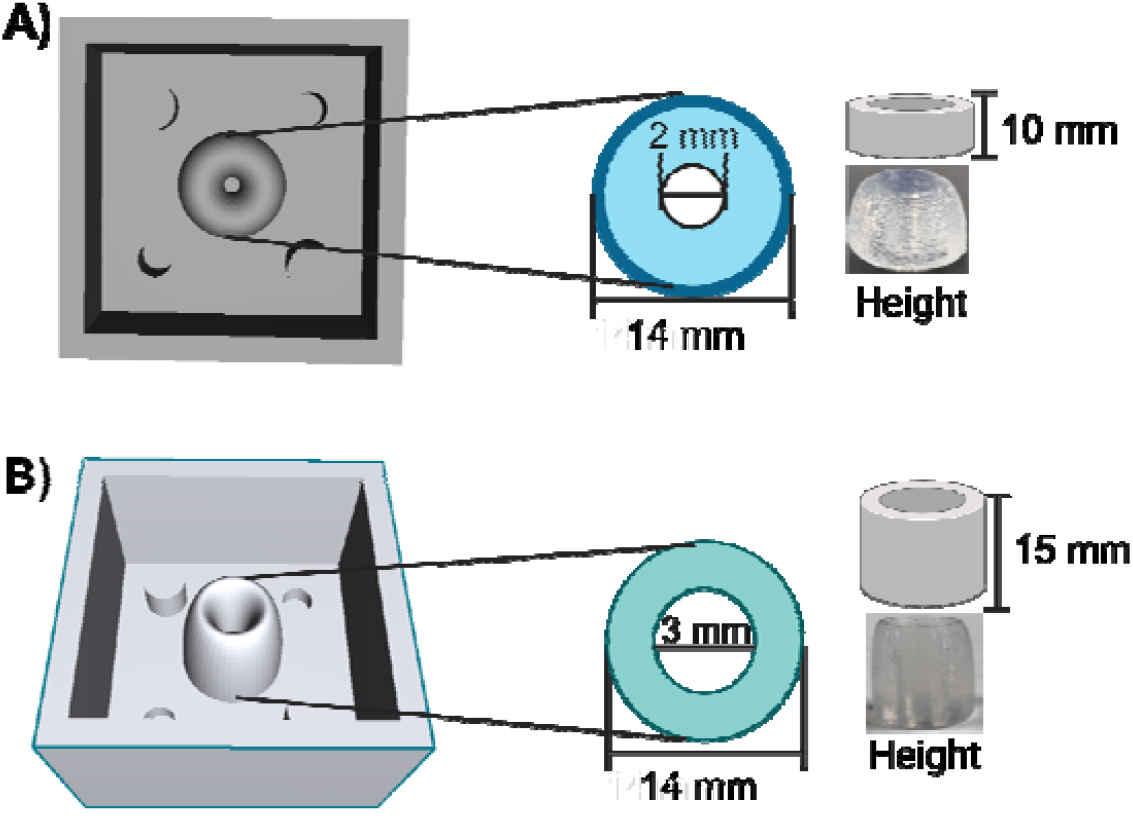
SolidWorks designed of the outside molds. (**A**) The first mold had a thickness of 10mm with an outside diameter of 14 mm and inside diameter of 2 mm. (**B**) The second mold had a thickness of 15 mm with an outside diameter of 14 mm and inside diameter of 3 mm. Images created in BioRender.com

**Supplementary Figure 2.**
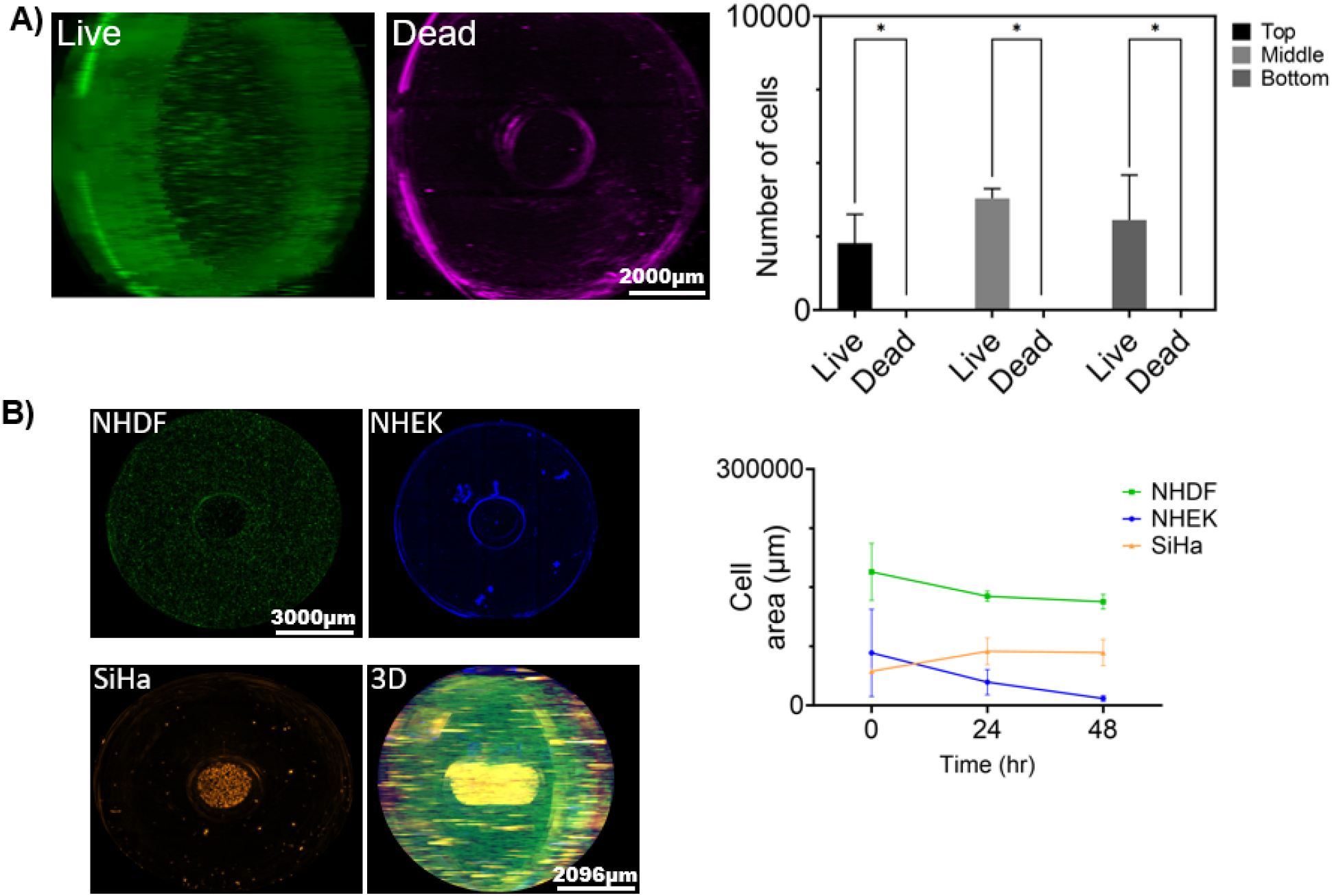
Cell response in 3D *in vitro* model with 15 mm thickness. **(A)** Cell viability and (**B**) Cell area were assessed in 3D *in vitro* model with 15 mm thickness. Representative images of live in green and dead cells in magenta after 48 hours of culture. The number of live and dead cells at 48 hours was quantified in the top, middle and bottom layers of the 3D model. (**B**) Representative images of the three cell lines (NHDF, NHEK, and SiHa) after 48 hours of co-culture in the 3D model. The cell area of each cell line was quantified at every time point. Images were taken every 24 hours for 48 hours. A montage of 4×5 (rows x columns) with 9 z-stacks was taken to capture the 3D model. The sample thickness was 1280 μm. Images were created using Gen5 V3.14 software, and post analysis was done in ImageJ. The data represent the mean ± SD (n = 4). \**p* < 0.05, analyzed using 2-Way ANOVA with Tukey post-test.

## Notes

### Competing Interest Statement

The authors have declared no competing interest.

